# Diversity in natural transformation frequencies and regulation across *Vibrio* species

**DOI:** 10.1101/683029

**Authors:** Chelsea A. Simpson, Ram Podicheti, Douglas B. Rusch, Ankur B. Dalia, Julia C. van Kessel

## Abstract

In marine *Vibrio* species, chitin-induced natural transformation enables bacteria to take up DNA from the external environment and integrate it into their genome via homologous recombination. Expression of the master competence regulator TfoX bypasses the need for chitin induction and drives expression of the genes required for competence in several *Vibrio* species. Here, we show that TfoX expression in two *Vibrio campbellii* strains, DS40M4 and NBRC 15631, enables high frequencies of natural transformation. Conversely, transformation was not achieved in the model quorum-sensing strain *V. campbellii* BB120 (previously classified as *Vibrio harveyi*). Surprisingly, we find that quorum sensing is not required for transformation in *V. campbellii* DS40M4. This result is in contrast to *Vibrio cholerae* that requires the quorum-sensing regulator HapR to activate the competence regulator QstR. However, similar to *V. cholerae*, QstR is necessary for transformation in DS40M4. To investigate the difference in transformation frequencies between BB120 and DS40M4, we used previously studied *V. cholerae* competence genes to inform a comparative genomics analysis coupled with transcriptomics. BB120 encodes homologs of all known competence genes, but most of these genes were not induced by ectopic expression of TfoX, which likely accounts for the non-functional natural transformation in this strain. Comparison of transformation frequencies among *Vibrio* species indicates a wide disparity among even closely related strains, with *Vibrio vulnificus* having the lowest functional transformation frequency. We show that ectopic expression of both TfoX and QstR is sufficient to produce a significant increase in transformation frequency in *Vibrio vulnificus.*

**Significance:** Naturally transformable or competent bacteria are able to take up DNA from their environment, a key method of horizontal gene transfer for acquisition of new DNA sequences. Our research shows that *Vibrio* species that inhabit marine environments exhibit a wide diversity in natural transformation capability ranging from non-transformable to high transformation rates in which 10% of cells measurably incorporate new DNA. We show that the role of regulatory systems controlling the expression of competence genes (*e.g.*, quorum sensing) is conserved among closely related species but differs throughout the genus. Expression of two key transcription factors, TfoX and QstR, are necessary and sufficient to stimulate high levels of transformation in *Vibrio campbellii* and recover low rates of transformation in *Vibrio vulnificus*.

## Introduction

Natural transformation is a physiological state in which bacteria are able to take up extracellular DNA from the environment, transport it across the cell envelope, and integrate it into their genome via homologous recombination. In marine *Vibrio* species, competence is induced by growth on the chitinous exoskeletons of crustaceans (1, 2). This process has been best studied in *Vibrio cholerae* (reviewed in (2)). Insoluble polysaccharide chitin is broken down by secreted extracellular chitinases (3). Soluble chitin oligosaccharides ultimately induce expression of the master competence regulator TfoX (4), which activates expression of numerous components of the competence machinery required to take up extracellular DNA. In addition to the chitin-sensing system, *V. cholerae* cells must also have a functional quorum-sensing system for natural transformation to be successful (1). Quorum sensing, the process of cell-cell communication, allows bacterial cells to respond to changes in population density and alter gene expression (5). The quorum-sensing systems in *Vibrio* species rely on detection of extracellular autoinducers that are sensed by membrane-bound sensor kinases, which shuttle phosphate to or away from the core response regulator LuxO at low cell density or high cell density, respectively (5). At the end of this phosphorylation cascade at high cell density, the master transcription factor called LuxR is activated, which controls expression of hundreds of genes (6). LuxR is the name of the master transcription factor in *Vibrio campbellii* (previously called *Vibrio harveyi*), whereas the homologs in other *Vibrio* species have different names: HapR (*V. cholerae*), SmcR (*Vibrio vulnificus*), and OpaR (*Vibrio parahaemolyticus*) (7). In *V. cholerae*, Δ*hapR* mutants are not competent, and this is due to HapR regulation of various competence genes, including *comEA, comEC, qstR*, and *dns* (1, 8, 9). HapR directly activates *qstR* (encoding a transcriptional regulator) and represses *dns* (encoding an extracellular DNase) (8). QstR subsequently activates downstream genes required for DNA uptake and integration, such as *comEA, comEC*, and *comM* (10). The requirement for HapR for competence can be circumvented if *qstR* and *tfoX* expression are induced and the *dns* gene is deleted (10). However, in wild-type *V. cholerae*, functional chitin-sensing and quorum-sensing systems are required for competence.

Growth on chitin is sufficient to induce competence in several *Vibrio* species, including *V. cholerae, V. vulnificus*, and *V. parahaemolyticus* (1, 11-13). Anecdotally, chitin-dependent natural transformation has not been observed in other species. In these cases, it may be that other environmental signals are required to induce competence or that laboratory conditions for chitin sensing are not sufficient. Further, multiple studies have identified different transformation capabilities among *V. cholerae* strains (14). However, because the chitin-sensing system drives expression of the core competence regulator TfoX, overexpression of Tfox is sufficient to bypass the requirement for chitin and induce competence in *V. cholerae, V. vulnificus, V. parahaemolyticus*, and *Vibrio natriegens* (1, 12, 15-17).

The type strain of *Vibrio campbellii* called BB120 (or ATCC BAA-1116) is a model system for studying quorum sensing in *Vibrio* species (18). This strain was historically called *Vibrio harveyi* until a recent comparative genomics analysis reclassified it as *V. campbellii (19)*. However, while BB120 is highly studied, the competence of this strain and others in this species was undetermined. Here, we show that the DS40M4 and NBRC 15631 environmental isolates of *V. campbellii* are highly competent following TfoX overexpression, whereas the lab strain BB120 and another environmental isolate called HY01 are not able to undergo natural transformation either via chitin induction or TfoX overexpression. We compare multiple *Vibrio* species and show that wide variation in transformation frequencies exists among strains in this genus.

## Results and Discussion

### *Induction of natural transformation via TfoX overexpression in* V. campbellii *DS4M04 and NBRC 15361 strains*

Anecdotal reports and unpublished experiments have shown that the *V. campbellii* type strain BB120 lacks the ability to undergo natural transformation. To formally test this, we assayed for transformation of plasmid DNA into BB120 using both chitin-dependent and - independent methods to induce competence (17, 20). For chitin-dependent transformations, cells are incubated with powdered chitin from shrimp shells and DNA and plated with antibiotic selection. For chitin-independent transformations, a plasmid expressing *V. cholerae tfoX* via an IPTG-inducible promoter (pMMB67EH-tfoX-kanR, Table S2) is used to induce competence in cells, and the cells are incubated with DNA prior to plating for antibiotic selection. When we performed both types of transformation experiments with BB120, we did not recover antibiotic-resistant colonies from either method, whereas high transformation frequencies were obtained with the same plasmid DNA introduced into positive control strains (*V. natriegens* for chitin-independent, Fig. S1A; *V. cholerae* for chitin-dependent, data not shown).

Because strains of *V. cholerae* exhibit differing natural transformation frequencies, we hypothesized that the absence of competence in BB120 may not be indicative of competence in other *V. campbellii* isolates. We therefore assayed natural transformation in three *V. campbellii* environmental isolates with sequenced genomes: DS40M4, NBRC 15631, and HY01 (19, 21-23). DS40M4 was isolated from the Atlantic Ocean near the west coast of Africa (24), HY01 was isolated from a bioluminescent shrimp in Thailand (22), and NBRC 15631 (also called CAIM 519 and ATCC 25920) was isolated from seawater in Hawaii (19). These species cluster together in the Harveyi clade in the *Vibrio* genus, with DS40M4 and NBRC 15631 being the most closely related (Fig. 1A). Incubation on chitin was unable to induce cells to take up the plasmid DNA in any of the *V. campbellii* strains (data not shown). However, both the DS40M4 and NBRC 15631 isolates were able to undergo transformation of plasmid DNA using the chitin-independent method in which TfoX was ectopically expressed (Fig. S1A). Transformation was dependent on TfoX expression because strains containing the empty vector control did not produce transformants (Fig. S1A).

**Figure 1.**
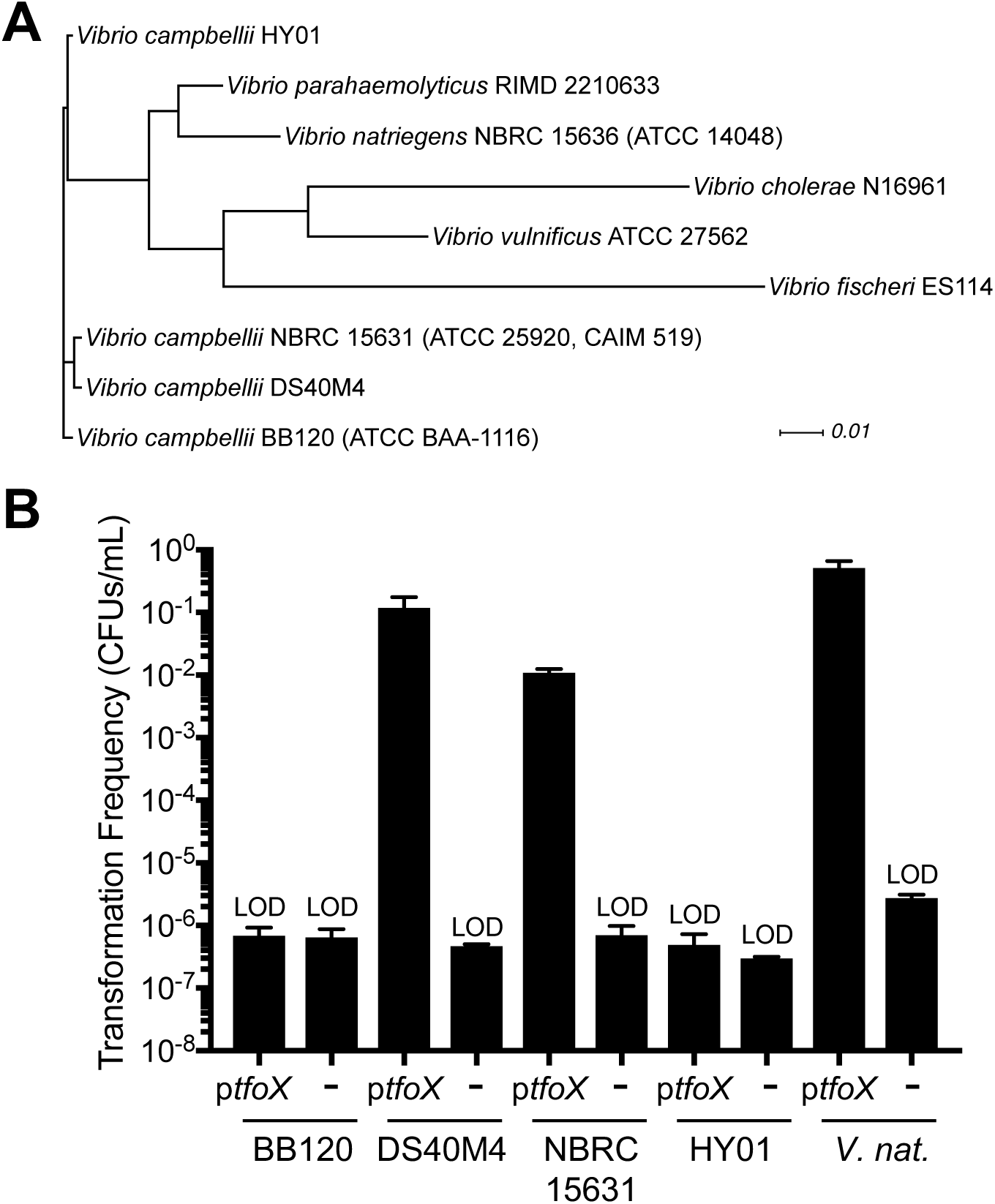
Natural transformation of *V. campbellii* strains via *tfoX* expression. (A) Phylogenetic tree of *Vibrio* strains based on comparison of amino acid sequences of 79 core conserved genes in the genomes shown. (B) Chitin-independent transformation of *V. campbellii* strains BB120, DS40M4, NBRC 15631, and HY01 compared to *V. natriegens*. Strains contained either a plasmid expressing *tfoX* (pMMB67EH-tfoX-kanR) or an empty vector control (pMMB67EH-kanR for *V. campbellii* strains or pMMB67EH-carbR for *V. natriegens*). Strains were transformed with 300 ng of linear *luxR::spec*^*R*^ tDNA (for *V. campbellii*) or *dns::spec*^*R*^ tDNA (for *V. natriegens*). LOD, limit of detection.

We next assayed transformation of linear DNA (referred to as transforming DNA, tDNA). We generated linear recombination products targeting the *luxO* gene in each *V. campbellii* strain containing an antibiotic resistance cassette and 3 kbp of DNA homologous to the regions flanking *luxO* for each strain (Δ*luxO::spec*^*R*^ substrates). High transformation frequencies were obtained for both DS4M04 and NBRC 15631 with their respective tDNAs that are dependent on the inducible expression of *tfoX* (Fig. 1B). We tested various parameters for transformation, including the number of cells in the reaction, the amount of time tDNA was incubated with cells before outgrowth, and the amount of tDNA added to the reaction (Fig. S1B, S1C, S1D). Using these optimized methods, we still did not observe transformation with either the BB120 strain or the HY01 strain (Fig. 1B). We note that the DS40M4 strain is close to or as efficient as *V. natriegens* at transformation with linear tDNA (Fig. 1B), which has thus far been reported as the *Vibrio* strain displaying the highest transformation frequencies (17). The lack of transformation observed in the highly pathogenic HY01 strain compared to the high frequencies observed in the oceanic isolate DS40M4 underscores a possible distinction in conservation of competence functions. As proposed in a recent study (10), it is possible that vibrios that are adapted to specific niches, such as those of a host organism for pathogenic strains, might have lost the ability to take up DNA if it were no longer beneficial. Conversely, natural transformation might be more advantageous for strains existing free-living in the ocean.

### *Assessment of quorum-sensing phenotypes in* V. campbellii *strains*

Because quorum sensing activates genes required for natural transformation in *V. cholerae*, we wanted to determine whether DS40M4 or NBRC 15631 contain functional quorum-sensing systems. Using chitin-independent natural transformation via *tfoX* induction, we generated Δ*luxO* and Δ*luxR* mutations in both the DS40M4 and NBRC 15631 strain backgrounds. In the model strain BB120, a Δ*luxO* mutant results in a constitutively expressed master quorum-sensing transcription factor LuxR, producing high levels of bioluminescence and mimicking a high cell density phenotype (25). Conversely, a Δ*luxR* mutant of BB120 is unable to produce bioluminescence (26). We compared the phenotypes of wild-type, Δ*luxO*, and Δ*luxR* strains for BB120, DS40M4, and NBRC 15361 by assessing expression of the quorum-sensing bioluminescence genes, *luxCDABE*. We were unable to monitor bioluminescence because DS40M4 and NBRC 15631 do not contain all of the *luxCDABE* bioluminescence genes like BB120 (they only encode *luxB* homologs) and do not bioluminesce (data not shown). We therefore used a GFP reporter plasmid in which the BB120 bioluminescence promoter (P_*luxCDABE*_) is transcriptionally fused to *gfp*. In BB120, a Δ*luxR* strain exhibits a 16-fold decrease in GFP expression compared to wild-type, whereas a Δ*luxO* mutant exhibits similar GFP levels as wild-type (Fig. 2A). In the DS40M4 strains, we observed similar GFP expression to the analogous BB120 strains (Fig. 2B). We complemented the Δ*luxR* deletion strain in DS40M4 and showed that GFP expression was restored to wild-type levels (Fig. S2A).

**Figure 2.**
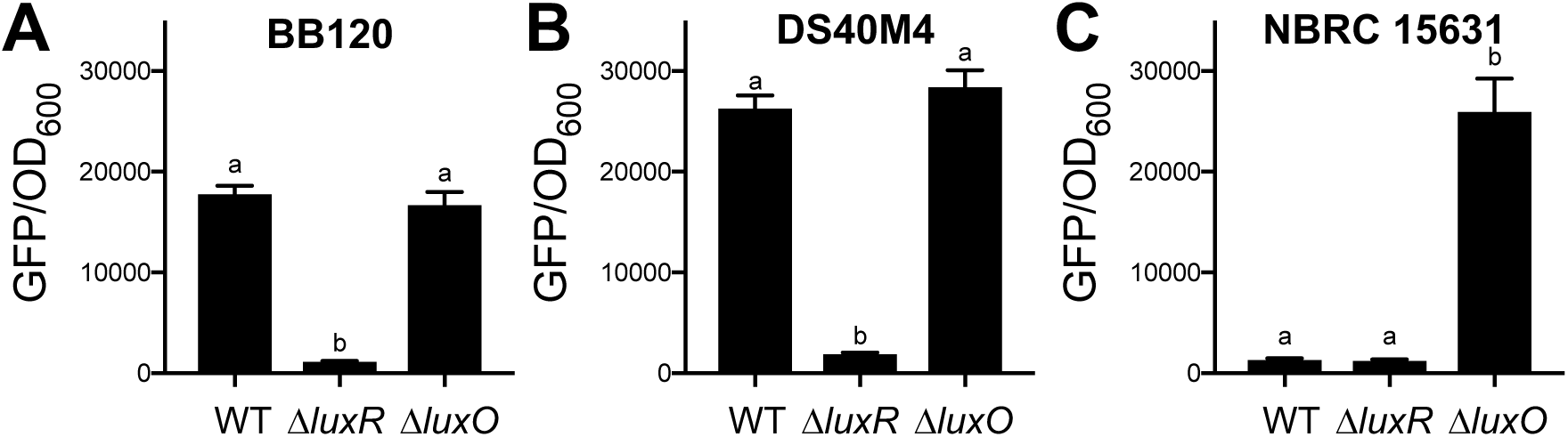
*V. campbellii* DS40M4 and NBRC 15631 encode functional LuxR and LuxO proteins. The P_*luxCDABE*_-*gfp* reporter plasmid pCS019 was introduced into wild-type, Δ*luxR*, and Δ*luxO* strains of *V. campbellii* BB120 (A), DS40M4 (B), and NBRC 15631 (C), and the GFP fluorescence divided by OD_600_ was determined. Different letters indicate significant differences (*p* < 0.001; one-way analysis of variation (ANOVA), followed by Tukey’s multiple comparison test of log-transformed data; *n* = 3).

The results with NBRC 15631 were different than BB120. The Δ*luxO* mutant exhibited higher levels of GFP compared to the Δ*luxR* mutant, as expected (Fig. 2C). Curiously, the wild-type NBRC 15361 strain consistently yielded low levels of GFP, even though the cells reached similar densities as the wild-type strains of BB120 and DS40M4 (Fig. 2C). This phenotype suggests that the NBRC 15631 cells do not produce and/or sense autoinducer(s) but retain functional *luxO* and *luxR* genes that are epistatic to autoinducer sensing. Strikingly, neither DS40M4 nor NBRC 15631 encode a LuxM homolog, which synthesizes AI-1 in BB120. DS40M4 and NBRC 15631 both encode close homologs of BB120 CqsS and LuxPQ, which bind CAI-1 and AI-2, respectively (98-99% amino acid identity). Conversely, LuxN, which binds AI-1, is less well conserved in DS40M4 and NBRC 15631 at 63-64% identity. There is high sequence conservation in the C-terminal receiver domain portion of LuxN, whereas the N-terminal region that binds autoinducers is not conserved with BB120. These data suggest that the DS40M4 and NBRC 15631 LuxN histidine kinases may recognize a different autoinducer than BB120. However, if this is the case, these two strains likely recognize the same molecule because the LuxN proteins are 99% conserved between DS40M4 and NBRC 15631. To determine if NBRC 15631 responds to an autoinducer from another strain, we incubated NBRC 15631 cells in media supplemented with cell-free supernatants from various *Vibrio* strains (including DS40M4). The GFP levels in the supernatant-treated cells were low and not significantly different than untreated NBRC 15631 cultures (data not shown). Importantly, we observed a high transformation frequency in NBRC 15631 Δ*luxO* that was similar to DS40M4. Thus, we suspect that there is a defect in regulation in the quorum-sensing pathway of NBRC 15631 that results in a low-cell-density-like state but can be bypassed by deletion of *luxO*. In this state, low levels of LuxR would be produced, which is likely the cause for the lower transformation frequencies in wild-type NBRC 15631 compared to DS40M4. From these data, we conclude that DS40M4 and NBRC 15631 have conserved LuxR and LuxO functions.

### *Quorum sensing is not required for transformation in* V. campbellii *DS40M4*

In *V. cholerae*, HapR expression is required for natural transformation (8, 27). In the absence of *hapR*, deletion of *dns* results in modest increases in transformation (28). A *hapR* mutation can be bypassed by over-expression of *qstR* to produce transformants, though at a reduced rate compared to wild-type (8). Combination of *qstR* expression and deletion of *dns* in a *hapR* mutant results in maximal transformation frequencies (10). To determine if the *luxR* genes of DS40M4 and NBRC 15631 are required for natural transformation, we assayed transformation frequencies in Δ*luxR* and Δ*luxO* mutants compared to wild-type for each *V. campbellii* strain using chitin-independent transformation. As a control, we performed the analogous experiment in *V. cholerae* with Δ*hapR* and Δ*luxO* mutants. Strikingly, we observed that a Δ*luxR* mutant in the DS40M4 strain is capable of natural transformation, though at a ∼60-fold reduced frequency compared to wild-type (Fig. 3A, 3B). The DS40M4 Δ*luxO* mutant exhibited a similar transformation frequency as wild-type DS40M4, which has also been observed in *V. cholerae* (1). This is due to maximal expression of LuxR/HapR in the Δ*luxO* background. The Δ*luxR* NBRC 15631 mutant did not yield colonies above the limit of detection (Fig. 3B). However, the Δ*luxO* NBRC 15631 strain had >100-fold higher levels of transformation than the wild-type NBRC 15631 strain and was similar to the frequencies observed in the DS40M4 Δ*luxO* strain (Fig. 3B). This result indicates that the presumed high levels of LuxR present in the Δ*luxO* strain restore transformation frequencies in NBRC 15631. From these data, we conclude that quorum-sensing control of competence via LuxR regulation is not required for transformation in *V. campbellii* DS40M4.

**Figure 3.**
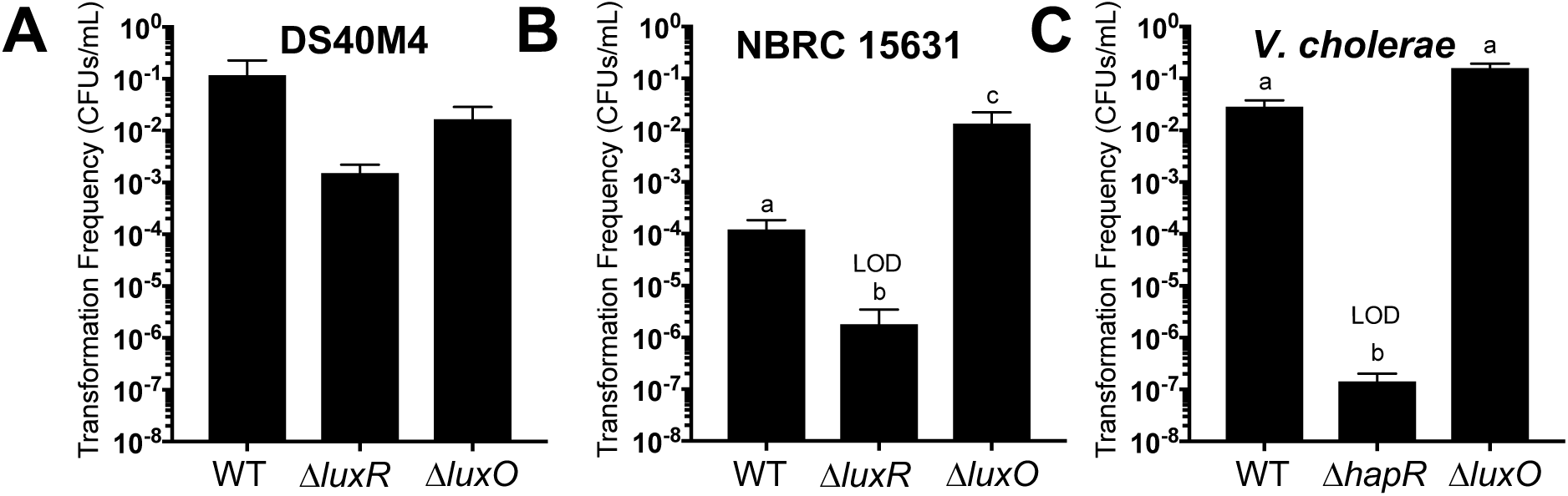
Natural transformation of *V. campbellii* DS40M4 does not require LuxR. (A, B) Transformation frequencies using chitin-independent transformation of a Δ*luxB::tm*^*R*^ substrate (300 ng) into wild-type, Δ*luxR*, and Δ*luxO* strains of DS40M4 (A) and NBCR 15631 (B). (C) Transformation frequencies using chitin-independent transformation of a Δ*vc1807::tm*^*R*^ substrate (300 ng) into wild-type, Δ*hapR*, and Δ*luxO* strains of *V. cholerae*. LOD, limit of detection. Different letters indicate significant differences (*p* < 0.05; one-way analysis of variation (ANOVA), followed by Tukey’s multiple comparison test of log-transformed data; *n* = 4).

### Conservation of competence genes and gene expression between BB120 and DS40M4

To investigate the differences in transformation frequencies between the BB120 and DS40M4 *V. campbellii* strains, we performed a comparative genomics analysis of the known *V. cholerae* competence genes against the genes present in *V. campbellii* DS40M4 and BB120. We generated a list of 47 *V. cholerae* genes based on published data supporting a role for the gene product and/or Tn-Seq data suggesting that mutants lacking these genes exhibit differing phenotypes in natural transformation assays. Our analyses show that BB120 and DS40M4 both encode homologs of all known genes that play a role in competence in *V. cholerae* (Fig. 4A, Table S4). Conservation of amino acid identity ranges from 25% to 95% with a median value of 73%. However, the vast majority of genes that have low amino acid conservation with *V. cholerae* are still highly conserved between BB120 and DS40M4. For only two genes, *pilA* and *VC0860* (a minor pilin gene), there is low conservation among the three strains. For example, the *pilA* gene in BB120 shares 44% amino acid identity with *V. cholerae pilA* and only 57% identity with DS40M4 *pilA* (Fig. 4A). PilA sequences vary widely among environmental *V. cholerae* strains (29), and it was recently shown that differences in PilA protein sequences among strains enable *V. cholerae* cells to discriminate between each other, which leads to decreased cell aggregation (30). Thus, it is unsurprising that the PilA sequence differs between all three vibrio strains (Fig. 4A). However, the VC0860 minor pilin protein is required for competence in *V. cholerae* (31), and thus the low conservation of this gene between BB120 and DS40M4 may reduce or eliminate transformation.

**Figure 4.**
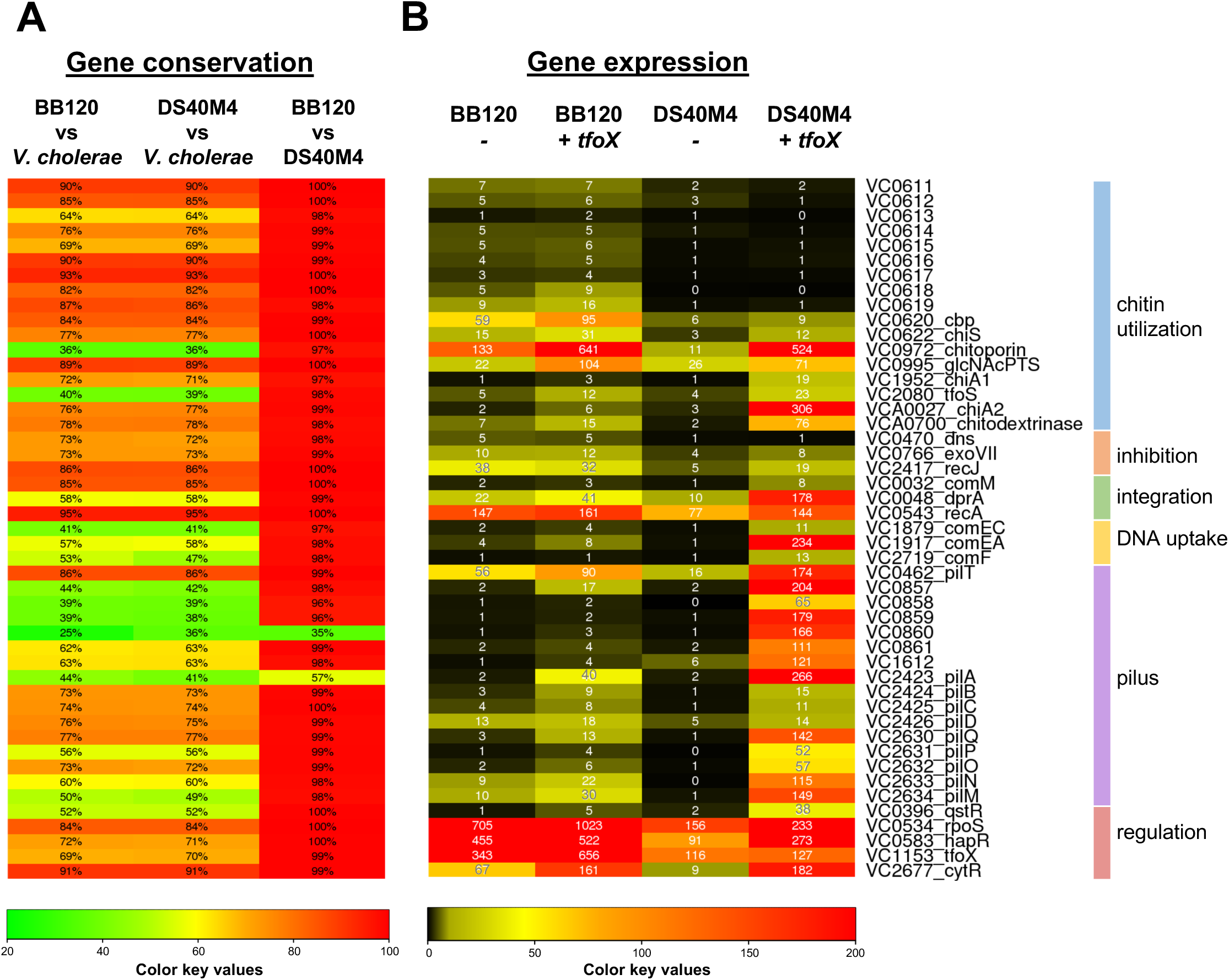
Comparative genomics and transcriptomics analyses of BB120 and DS40M4 competence genes. Genes required for DNA uptake and integration previously determined in *V. cholerae* were identified in BB120 and DS40M4 using reciprocal BLAST analyses. Genes are organized based on function. The locus tags correspond to *V. cholerae* genes; corresponding locus tags for BB120 and DS40M4 are in Table S4. (A) The chart indicates the amino acid identity shared between *V. cholerae*, BB120, and DS40M4, which is shown in each bar and by the color scale. (B) The chart indicates the RPKM values derived from RNA-seq data comparing either BB120 or DS40M4 transcript levels in the presence or absence of *tfoX* induction (strains contain plasmid pMMB67EH-tfoX-kanR).

Because it appears that BB120 encodes all known genes required for competence at some degree of conservation, we next questioned whether the expression of these genes was sufficient for transformation. We performed RNA-seq comparing gene expression in the presence or absence of *tfoX* induction in both BB120 and DS40M4 strains and analyzed the reads per kilobase of transcript per million mapped reads (RPKM). As expected, induction of TfoX expression resulted in upregulation of the genes required for DNA uptake and integration (*comEC, comEA, comM, comF*, and *dprA*) and pilus structure (*pilABCD, pilMNOPQ*, minor pilins VC0857-0861) in DS40M4 (Fig. 4B). However, for most of these genes in BB120, there is no significant change in expression +/- *tfoX* induction (Fig. 4B). Further, even if an increase in gene expression was observed in BB120 with *tfoX* induction, it was modest compared to the large changes in gene expression in DS40M4 with *tfoX* induction. Collectively, these data suggest that expression levels of competence genes in BB120 are not sufficient for transformation. It is also possible that the *pilA* and *VC0860* homologs in BB120 are not functional, due to their lower conservation with DS40M4 homologs.

Over-expression of *V. cholerae tfoX* in *V. campbellii* DS40M4 produced a similar expression profile for the 47 competence genes to the profile produced in *V. cholerae* upon *tfoX* induction (32). One exception to this is that we observed a ∼10-fold increase in *pilT* expression with *tfoX* induction, as compared to *V. cholerae* that showed a similar level of *pilT* expression in the presence or absence of *tfoX* induction (32). Another exception is that *tfoX* induction in DS40M4 resulted in a ∼20-fold increase in *cytR* expression in DS40M4, whereas in a similar experiment in *V. cholerae, cytR* expression is only increased ∼4-fold (32). Although the reason for these differences is unclear, it minimally indicates that there are slight differences in the TfoX regulons of *V. campbellii* DSS40M4 and *V. cholerae*. Overall, our observation that DS40M4 up-regulates >20 genes similar to *V. cholerae* in the presence of TfoX suggests high conservation of the regulatory network controlling competence.

We noted that the fold-changes in the *qstR* and *cytR* transcript levels are larger in DS40M4 +/- *tfoX* induction as opposed to BB120 +/- *tfoX* induction. This is important because both QstR and CytR are transcriptional regulators; CytR is a transcription factor that positively regulates competence genes, among others, in *V. cholerae* (33). The levels of *cytR* in BB120 with *tfoX* induction are similar to those in DS40M4 with *tfoX* induction, suggesting that low CytR expression is not likely the cause for lack of transformation in BB120 (Fig. 4B). Conversely, the levels of *qstR* in BB120 are lower than in DS40M4 under *tfoX* induction conditions (Fig. 4B). Though we cannot compare RPKM values for genes between strains, by normalizing to other genes such as *recA* of each strain, we can see that the levels of *qstR* are lower in BB120 (Fig. 4B). Because *qstR* is required for expression of essential competence genes (*e.g., comEA, comEC*) in *V. cholerae* (10), the low levels of *qstR* expression in BB120 might be one reason for its inability to take up DNA.

### Expression of QstR in DS40M4 drives maximum levels of natural transformation in the absence of LuxR

We next sought to examine the mechanism behind the observation that the Δ*luxR* strain of DS40M4 retains transformation capability. We hypothesized that this may be due to either 1) higher levels of *qstR* expression in DS40M4 than in *V. cholerae* (in the absence of LuxR/HapR), or 2) poorly functional or poorly expressed *dns* in DS40M4 compared to *V. cholerae*. To formally test whether increased levels of *qstR* would increase transformation frequencies, we cloned the *qstR* gene from DS40M4 downstream of *tfoX* under control of an IPTG-inducible promoter to form a synthetic operon: P_*tac*_*-tfoX-qstR* (pCS32). We observed a ∼20-fold increase in transformation frequency in DS40M4 Δ*luxR* with the expression of *qstR*, which brought the level in this background to the wild-type frequency (Fig. 5A), suggesting that Dns activity may not play a role in transformation frequency as seen in *V. cholerae*. Deletion of *qstR* in DS40M4 abolishes transformation, much like in *V. cholerae* (Fig. 5B). These data indicate that QstR expression is epistatic to LuxR. From these data, we conclude that TfoX and QstR are both necessary and sufficient to initiate natural transformation in *V. campbellii* DS40M4 in the absence of LuxR. Unfortunately, we did not observe any transformation in BB120 or HY01 under induction of both *tfoX* and *qstR* (data not shown).

**Figure 5.**
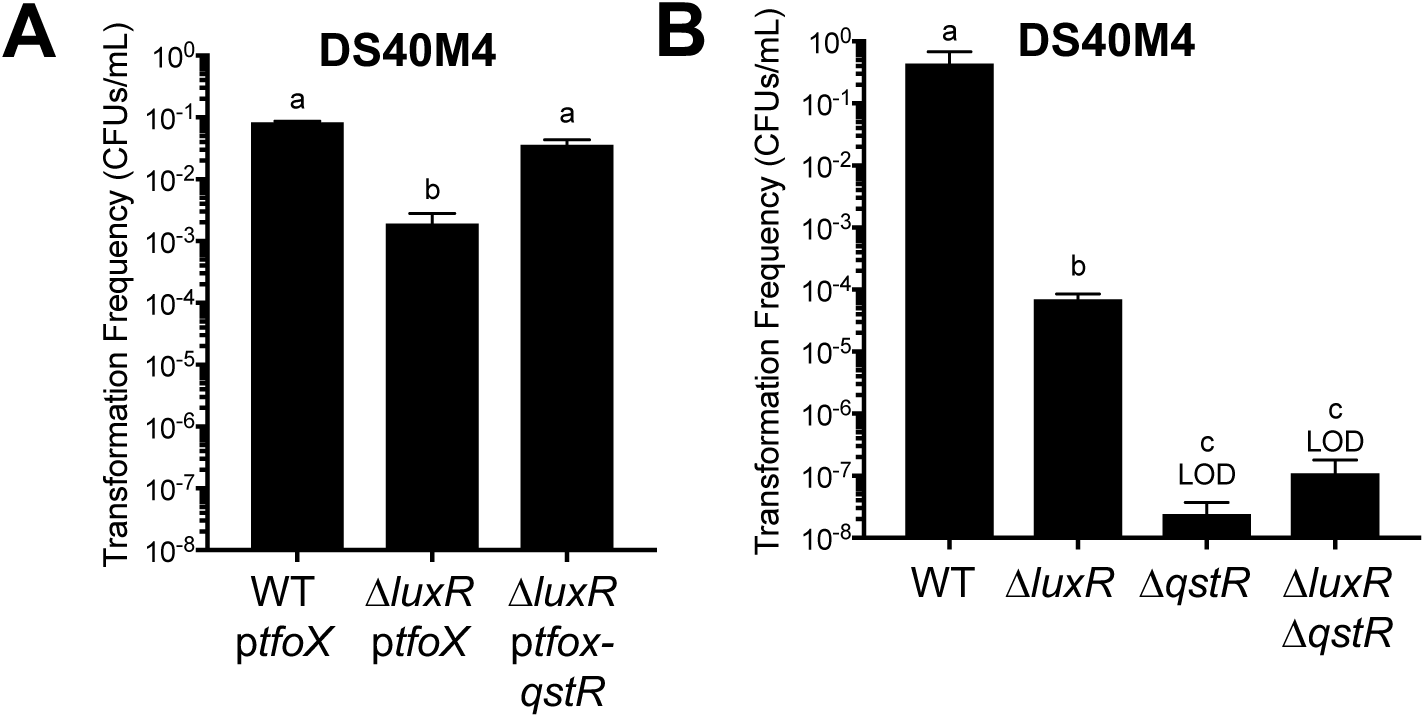
QstR and TfoX are necessary and sufficient to bypass LuxR for natural transformation. (A) Transformation frequencies using chitin-independent transformation of a Δ*luxB::tm*^*R*^ substrate (300 ng) into wild-type and Δ*luxR* strains of DS40M4, either with the pMMB67EH-tfoX-kanR plasmid (p*tfoX*) or pCS32 plasmid (p*tfoX-qstR*). (B) Transformation frequencies using chitin-independent transformation of a Δ*luxB::spec*^*R*^ substrate (300 ng) into wild-type, Δ*luxR*, Δ*qstR*, and Δ*luxR* Δ*qstR* strains of DS40M4, all containing pMMB67EH-tfoX-kanR. Different letters indicate significant differences (*p* < 0.01; one-way analysis of variation (ANOVA), followed by Tukey’s multiple comparison test of log-transformed data; *n* = 3).

### *Natural transformation frequencies vary widely across* Vibrio *species*

The notable differences in transformation frequencies between strains of *V. campbellii* led us to investigate the frequencies of natural transformation in other vibrios. Varied transformation frequencies have been reported for other *Vibrio* species in the literature, with *V. natriegens* having the highest level of transformation frequency (10, 12, 16, 17, 34). However, several of these experiments were performed in different labs with different conditions, such as different *tfoX* genes, tDNA homology lengths, tDNA quantity, media, and outgrowth time. To formally assay frequencies of transformation with consistent experimental conditions, we tested transformation using both chitin-dependent and –independent methods in multiple *Vibrio* species: *V. campbellii, V. parahaemolyticus, V. cholerae*, and *V. vulnificus*. The tDNA substrates were generated to target the gene encoding the LuxR/HapR homolog in each strain: *luxR, opaR, hapR*, and *smcR*, respectively. The only difference in experimental conditions was the use of media containing higher concentrations of salt required for viability in *V. campbellii, V. vulnificus*, and *V. parahaemolyticus*. In our experiment, only *V. cholerae* was able to undergo natural transformation when chitin was used to induce competence (Fig. S2B). This is in contrast to other studies that have successfully used crab shell pieces to induce transformation in *V. vulnificus* and *V. parahaemolyticus* (12, 13). However, another group was unable to induce competence in *V. parahaemolyticus* with long-chain chitin polymers (16). Thus, it is important to note that we only used chitin from shrimp shells and did not test the effect of soluble chitin oligosaccharides or crab shells, which may produce better transformation frequencies.

When we stimulated competence by overexpressing TfoX (chitin-independent), all strains except *V. campbellii* BB120 and HY01 were able to take up the tDNA and produce recombinants (Fig. 6A). *V. vulnificus* exhibited the lowest transformation frequency, and *V. natriegens* and *V. campbellii* DS40M4 have the highest frequencies (Fig. 1B, Fig. 6A). Similar to a previously published comparison, *V. parahaemolyticus* had a similar transformation frequency to *V. cholerae* (35). Because expression of both *tfoX* and *qstR* were sufficient to increase transformation frequency in DS40M4 in the absence of *luxR*, we hypothesized that expression of these two transcription factors might increase transformation frequency in other vibrios. We introduced the same P_*tac*_*-tfoX-qstR* plasmid pCS32 into each *Vibrio* strain tested above and assayed for transformation frequency. We observed a >2-log increase in transformation in *V. vulnificus* (Fig. 6A), though no significant increases in the other vibrios tested. We suspect that ectopic expression of *qstR* is likely compensating for low levels of *qstR* expression in *V. vulnificus*. Of note, this is a drastic improvement of transformation frequency in this species, which is otherwise too low for multiplex genome editing by natural transformation (MuGENT) (34). Our observed variations in transformation frequencies across *Vibrio* species may be due to either differences in expression and/or function of Dns and/or varying levels of QstR in different strains.

**Figure 6.**
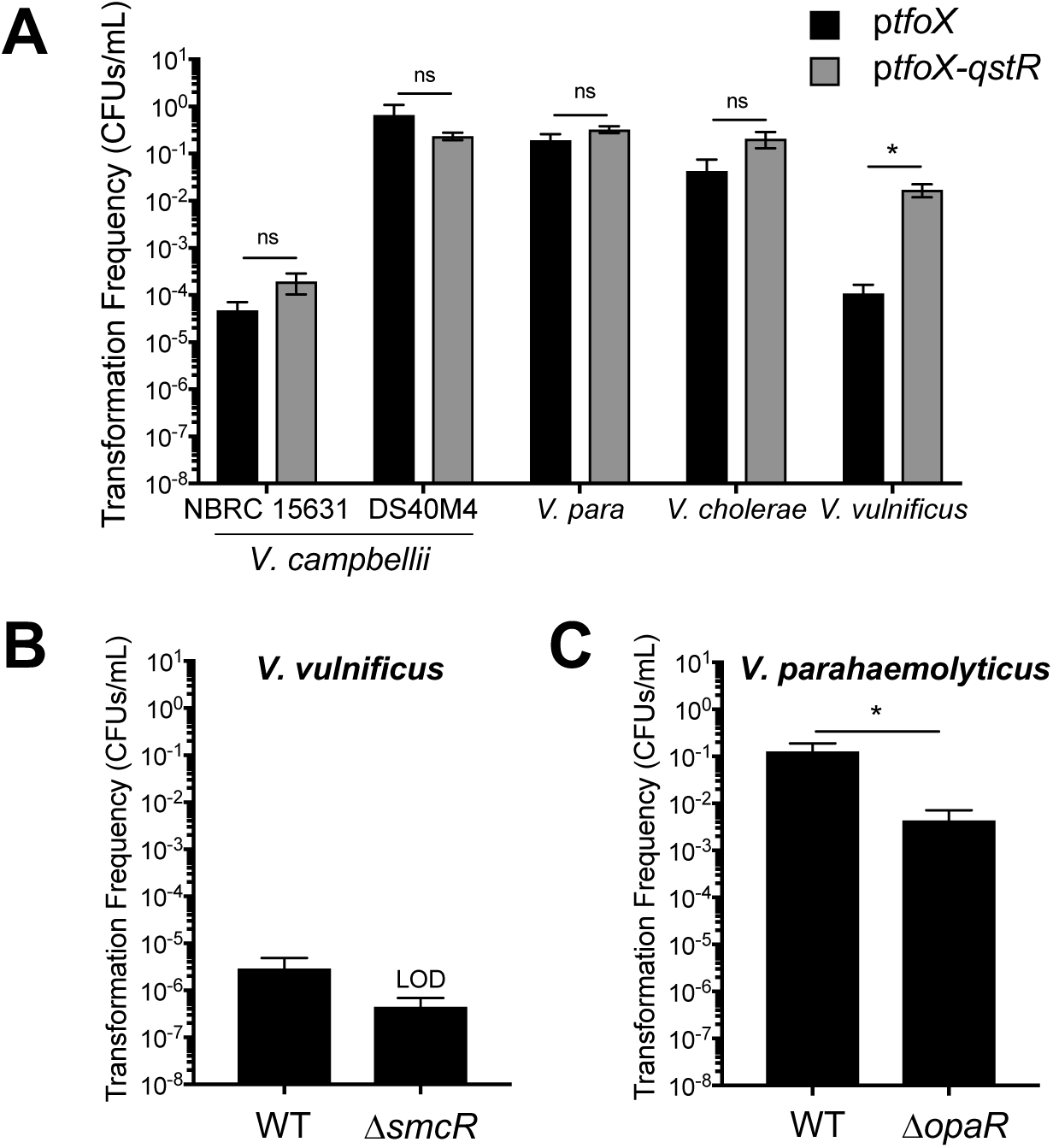
Natural transformation frequencies vary in *Vibrio* species. (A) Chitin-independent transformations in each of the listed *Vibrio* species using Spec^R^ linear tDNAs (300ng) targeting the *luxR* homolog in each strain in the presence of a plasmid expressing either *tfoX* alone (pMMB67EH-tfoX-kanR) or both *tfoX* and *qstR* (pCS32). Asterisks indicate significant differences (*p* < 0.0001; two-way analysis of variation (ANOVA), followed by Sidak’s multiple comparison test of log-transformed data; *n* = 3). (B, C) Transformation frequencies using chitin-independent transformation of a Δ*pomB::tm*^*R*^ substrate (300 ng) into wild-type and Δ*smcR V. vulnificus* strains expressing *tfoX* (pMMB67EH-tfoX-kanR) (B) or wild-type and Δ*opaR V. parahaemolyticus* strains expressing *tfoX* (pMMB67EH-tfoX-kanR) (C). The asterisk indicates *p* < 0.05 in an unpaired, two-tailed t-test.

### *Quorum sensing is not required for transformation in* V. parahaemolyticus

To determine if the lack of requirement for LuxR for competence extends beyond *V. campbellii*, we tested transformation in *V. vulnificus* and *V. parahaemolyticus* in the presence or absence of the LuxR-type quorum-sensing regulator in each species. Deletion of the *smcR* gene in *V. vulnificus* abolished transformation, similar to what is observed in *V. cholerae* (Fig. 6B). However, deletion of the *opaR* gene in *V. parahaemolyticus* did not eliminate transformation, but decreased it ∼30-fold, similar to what is observed in *V. campbellii* DS40M4 (Fig. 6C, Fig. 3A). Minimally, these data show that the regulatory factors (*e.g.*, QstR, LuxR/HapR) controlling transformation are conserved in *V. cholerae, V. parahaemolyticus*, and *V. campbellii* DS40M4, but the stringency of dependence on these factors varies among species. These differences may also be reflective of their evolutionary relatedness; *V. parahaemolyticus* and *V. campbellii* are more closely related to each other than to *V. cholerae* and *V. vulnificus* (Fig. 1A). It may also reflect niche-specific requirements for DNA uptake that differ in free-living strains compared to host-associated strains.

## Concluding Remarks

In this study, we identified two strains of *V. campbellii* that are capable of natural transformation in the presence of *tfoX* expression. We observed a wide diversity in transformation frequencies, not only among *campbellii* strains but across *Vibrio* species, some which are highly competent and some that are incapable of transformation under tested laboratory conditions. Our results show that the connection between quorum sensing and competence is linked but not dependent in some *Vibrio* strains. Unfortunately, we were unsuccessful at restoring competence to *V. campbellii* BB120. It is likely that the limitations to transformation in BB120 are multifactorial, including a potentially defective PilA and/or VC0860 minor pilin homolog, decreased gene expression of genes required for pilus assembly, DNA uptake, and DNA integration upon induction of *tfoX*, and low expression of QstR. It is also not clear why so many competence genes failed to induce upon TfoX expression, whereas this method of inducing competence was highly successful in *V. campbellii* DS40M4 and NBRC 15631 and other *Vibrio* species. We postulate that either *V. cholerae* TfoX does not function properly in BB120 or that BB120 lacks regulatory control of the core competence genes from another transcription factor, possibly one that is not known in *V. cholerae* yet. The only BB120 gene known to be controlled by TfoX that responded with its induction was *qstR*, suggesting that TfoX might function in BB120 but perhaps at a reduced capability. This is in accordance with the finding that the *V. fischeri tfoX* is slightly decreased in the ability to cross-activate the *qstR* of *V. cholerae* (35). One additional possibility is that there is a regulatory feedback loop that downregulates transcription levels of structural genes in the absence of a functional pilus (*i.e.*, if the BB120 PilA is not functional), similar to the homeostatic regulation of flagellin in *Bacillus* (36). Future experiments are required to dissect the broken pieces of the BB120 competence regulatory and functional networks to synthetically generate competence in this model organism.

## Materials and Methods

### Bacterial Strains and Media

All bacterial strains and plasmids used are listed in Table S1. When necessary, media was supplemented with carbenicillin (100 μg/ml), kanamycin (50 µg/ml for *E. coli*, 100 μg/ml for vibrios), spectinomycin (200 µg/ml), chloramphenicol (10 μg/ml), and/or trimethoprim (10 µg/ml). Plasmids were transferred from *E. coli* to *Vibrio* strains by conjugation on LB plates. Exconjugants were selected on LB or LM plates with polymyxin B at 50 U/ml and the appropriate selective antibiotic. For descriptions of growth media used for each strain, see SI Materials and Methods.

### Molecular Methods

All PCR products were synthesized using Phusion HF polymerase (New England Biolabs). Sequencing of constructs and strains was performed at Eurofins Scientific. Cloning procedures and related protocols are available upon request. Agarose gels (0.8-2%) were used to resolve DNA products. Oligonucleotides used in the study are listed in Table S3.

### Synthesis of Linear tDNAs

Linear tDNAs were generated by splicing-by-overlap extension (SOE) PCR as previously described by Dalia *et al*. (15). For details, see SI Materials and Methods.

### Natural Transformation

Chitin-independent transformations were performed following the protocol established in Dalia *et al.* (17). Chitin-dependent transformations were performed following the protocol established in Dalia *et al.* (20). For details, see SI Materials and Methods. Following natural transformation, strains containing the correct target mutation were identified via colony PCR with a forward and reverse detection primer (Table S3).

### GFP assays

*Vibrio* strains were first cured of the pMMB67EH-tfoX-kanR plasmid by serial inoculation (3-4 times) in the absence of antibiotic selection. The pCS19 reporter plasmid was conjugated into strains. Overnight cultures were diluted 1:5,000 into fresh media and incubated at 30°C shaking overnight. GFP fluorescence and OD_600_ were measured in black 96-well plates using the BioTek Cytation 3 Plate Reader.

### RNA-seq

Strains were inoculated in 5 ml LBv2 and grown overnight shaking at 250 RPM at 30°C. Each strain was back-diluted to an OD_600_= 0.005 in fresh LBv2 (uninduced control samples) or LBv2 with 100 μM IPTG (induced samples). Cultures were grown shaking at 250 RPM at 30°C until they reached OD_600_ =∼1.8. Cells were collected by centrifugation at 3700 RPM at 4°C for 10 min. The supernatant was removed, the pellets flash frozen in liquid N_2_, and stored at −80°C. RNA was isolated using the Qiagen RNeasy Mini Kit following the manufacturers protocol and treated with DNase (Ambion). RNA-seq was performed at the Center for Genomics and Bioinformatics at Indiana University. The results were deposited in the National Center for Biotechnology Information Gene Expression Omnibus database (NCBI GEO). For details on RNA sequencing and data analyses, see SI Materials and Methods.

### Comparative Genomics and Phylogenetics

Reciprocal BLASTs between DS40M4 and BB120 were performed using NCBI BLASTP ver. 2.7.1. following re-annotation of Genbank sequences with Prokka ver. 1.12 (37). To generate the phylogenetic tree, a multiple sequence alignment was performed with Genbank sequences using MUSCLE ver. 3.8.31 (38), input into the program RAxML ver. 8.2.12 (39), and a best scoring maximum likelihood tree was constructed. For details, see SI Materials and Methods.

## Acknowledgments

We thank Triana Dalia for guidance on SOE PCR. We also thank Evan Schneider for technical assistance. We thank Margo Haywood for the DS40M4 strain and Varaporn Vuddhakul for the HY01 strain. This work was supported by National Institutes of Health grants R35GM124698 to JVK and R35GM128674 to ABD.

