## Supplemental Information for "Diversity in natural transformation frequencies and regulation across *Vibrio* species"

Supplemental Materials and Methods
Supplemental Figures (Figures S1-S2)
Supplemental Tables (Tables S1-S4)
Supplemental References

### Supplemental Materials and Methods

*Bacterial Strains and Media*

*E. coli* and *V. cholerae* were grown in Lysogeny broth (LB) for all experiments. *V. parahaemolyticus* was grown either on LBv2 (LB medium supplemented with additional 200 mM NaCl, 23.14 mM $MgCl_2$, and 4.2 mM KCl) for chitin-independent transformations or on LMv3 (LB medium supplemented with additional 513 mM NaCl) for chitin-dependent transformations. *V. campbellii*, *V. natriegens*, and *V. vulnificus* were grown on Luria Marine (LM) medium for chitin-dependent transformations (LB medium supplemented with additional 171 mM NaCl) or LBv2 for chitin-independent transformations. Instant ocean water (IOW) medium was used in the chitin-dependent and independent transformations, which is Instant Ocean Sea Salts (Aquarium Systems, Inc.) diluted in sterile water (0.5X = 7 g/l, 2X = 28 g/l). IOW at 0.5X was used for *V. cholerae*, and IOW at 2X was used for all other strains. Chitin slurry consists of 8 g of chitin powder from shrimp shells (Sigma-Aldrich) in 150 ml of 0.5X IOW.

*Synthesis of Linear tDNAs*

Three-piece splicing-by-overlap extension (SOE) transforming DNAs (tDNAs) were synthesized in which the target gene was replaced with an antibiotic-resistance cassette. The three pieces of the tDNA consisted of 1) an UP arm which contained ~3kb upstream region of homology amplified with F1 and R1 primers, 2) an antibiotic-resistance cassette ($spec^R$, $TM^R$), and 3) a DOWN arm which contained ~3kb downstream homology amplified with F2 and R2 primers (refer to F1, R1, F2, R2 names in Table S3). The segments were gel purified, mixed in equal ratios, and used as templates in a PCR reaction where the construct was stitched together using the F1 and R2 primers.

*Natural Transformation*

All experiments used pMMB67EH-kanR (empty vector control), pMMB67EH-tfoX-kanR (*tfoX* expression plasmid), or pCS32 (*tfoX-qstR* expression plasmid), with the only exception being the use of pMMB67EH-carbR as an empty vector control in *V. natriegens*. Strains containing *tfoX* expression plasmids were grown overnight (12-18 hr) in LBv2 or LB with antibiotic selection and 100 μM IPTG shaking at 250 RPM at 30°C to induce competence. Subsequently, 7 μl of the overnight culture was diluted directly into 350 μl of either 0.5X IOW (*V. cholerae*) or 2X IOW (all other vibrios) supplemented with 100 μM IPTG. Transforming DNA was added as indicated for each experiment, and the reactions were incubated statically at 30°C for 4-5 hr. One ml of LB (*V. cholerae*) or LBv2 (all other vibrios) was added to each reaction, and these were grown at 30°C shaking at 250 RPM for ~1-2 hrs. Reactions were plated on selective media to select for integration of the tDNA and on non-selective media to determine total viable cells. Transformation frequency was calculated as the number of antibiotic-resistant colonies divided by viable cells.

Cultures were grown overnight in LB, LM, or LMv3 (depending on the *Vibrio* species as described above) shaking at 250 RPM at 30°C. A subculture was made by diluting 13 μl of the overnight culture into 3 ml fresh LM or LB and growing shaking at 30°C until the $OD_{600}$ = 0.5-1.0. Subsequently, 1 ml of cells was pelleted at 13, 000 RPM for 1 min, the supernatant was removed, and the cells were resuspended to an $OD_{600}$=1.0 in 0.5X or 2X IOW (depending on the *Vibrio* species as described above). Next, 100 μl of the resuspended cells was added to a 2-ml tube containing 150 μl chitin slurry and 750 μl 0.5X or 2X IOW. The reaction was vortexed briefly and incubated statically at 30°C for 16-24 h. Approximately 550 μl of supernatant was removed from each reaction without disturbing the settled chitin. The tDNA was added (quantities indicated in figure legends), the tube inverted gently to mix, and the reaction allowed to incubate statically at 30°C for 16-24 h. LM or LB (0.5 ml) was added, and reactions were outgrown at 30°C for 3-5 h before plating on selective and non-selective media.

*RNA-seq*

Isolated RNA was treated with Ribo-Zero rRNA (Bacteria) removal kit (Illumina). Purified RNA was prepared using TruSeq Stranded mRNA HT Sample Prep kit (Illumina) according to manufacturer's protocols; dual-indexed adapters were added to libraries for multiplexing and then libraries were cleaned by AMPure XP beads (Beckman Coulter).

Sequencing reads were trimmed using Trimmomatic (version 0.38; (1)) with a minimum trimmed read length of 30. The trimmed reads were mapped on to the Vibrio campbellii BB120 genome using bowtie2 [version 2.3.4.3 with default parameters; (2)]. Read counts for genes and intergenic intervals were calculated using a custom perl script. Resulting gene/interval counts were used to conduct differential expression analysis using the program DESeq2 algorithm with default parameters (3).

*Reciprocal BLAST*

Genome sequences and annotated protein sequences for *V. campbellii* strains BB120 (accessions CP000789.1, CP000790.1, CP000791.1) and DS40M4 (accessions CP030788.1, CP030789.1, CP030790.1) were obtained from Genbank. To ensure that we did not miss any unannotated genes, the genome sequences were re-annotated with Prokka ver. 1.12 (4) (parameters: --minpid 70 --usegenus --hmmlist TIGRFAM, CLUSTERS, Pfam, HAMAP). Protein sequences encoded by annotated genes from the following strains were used as the training set for Prokka predictions: *V. campbellii* strains BB120 (ATCC BAA-1116), DS40M4, HY01, and NBRC 15631 (ATCC 25920, CAIM 519), *V. cholerae* N16961, *V. fischeri* ES114, *V. natriegens* NBRC 15636 (ATCC 14048), *V. parahaemolyticus* RIMD 2210633, and *V. vulnificus* ATCC 27562. Protein sequence sets for each of the two *V. campbellii* strains BB120 and DS40M4 were prepared by combining sequences from Genbank annotation with those from Prokka predictions. The two protein sequence sets were aligned against each other using NCBI BLASTP ver. 2.7.1. For all the sequences in each set, the best scoring matches in the other set were identified based on the highest bit score. When multiple best hits were encountered, the Genbank annotated gene maintaining synteny with the best hits for the neighboring genes was preferred over the rest. Orthologous gene pairs for the two strains were called when the protein sequences encoded by the two genes identified each other as their best match.

*Phylogenetic construction*

Nucleotide sequences from the coding regions for the annotated genes were downloaded from Genbank for the nine *Vibrio* species. The sequences from *V. campbellii* BB120 were aligned against those from the remaining eight genomes. The core genome was defined by identifying genes present in all nine *Vibrio* strains with at least 70% DNA identity over at least 80% of the total length of the corresponding homologous gene in BB120 using the method as described (5). This formed the core set of 79 genes specific to these nine *Vibrio* strains. The protein sequences for the core gene set across all the nine genomes were obtained, and a multiple sequence alignment was performed separately for each gene using MUSCLE ver. 3.8.31 (6). After trimming the ends of the alignments to remove contiguous gaps (if any) resulting from unequal gene lengths, the alignments from each gene for each species were concatenated in order. Using this combined amino acid multiple sequence alignment file as input to the program RAxML ver. 8.2.12 (7), a best scoring maximum likelihood tree was constructed.

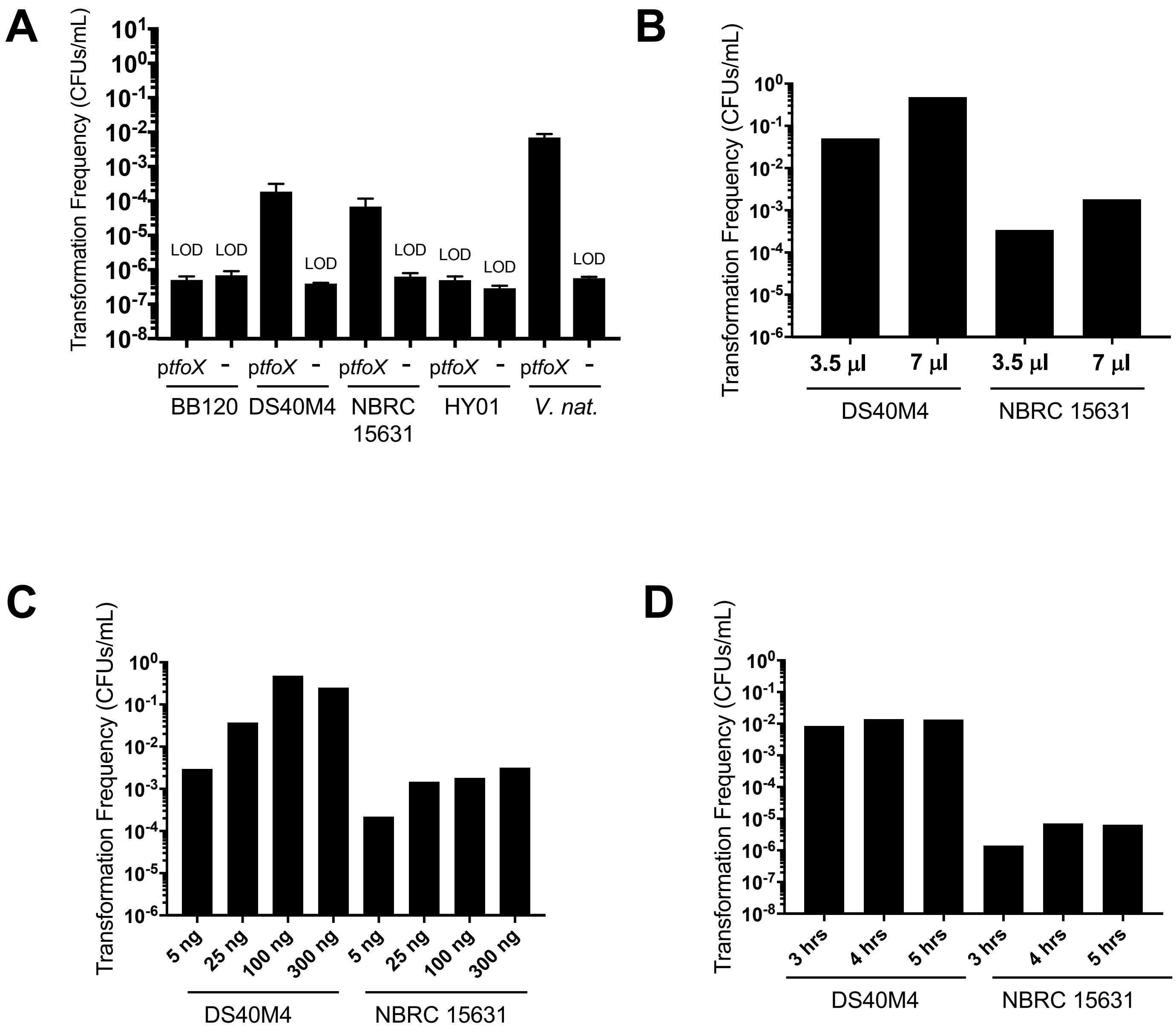


**Figure S1.** (A) Chitin-independent transformations of *V. campbellii* strains BB120, DS40M4, NBRC 15631, and HY01 with plasmid tDNA compared to positive control strain *V. natriegens.* Strains contained either a plasmid expressing *tfoX* (pMMB67EH-tfoX-kanR for *V. campbellii* strains) or an empty vector control (pMMB67EH-kanR for *V. campbellii* strains or pMMB67EH for *V. natriegens*). Strains were transformed with 1 µg of plasmid tDNA conferring chloramphenicol resistance (pJV298). LOD, limit of detection. (B) Chitin-independent transformation efficiencies in DS40M4 and NBRC 15631 reactions with varied quantities of cell inoculations. Strains were transformed with 50 ng linear *luxO::spec*$^R$ tDNA. Averaged results from two technical replicates are shown for each condition. (C) Chitin-independent transformation efficiencies in DS40M4 and NBRC 15631 reactions with varied quantities of linear *luxO::spec*$^R$ tDNA. Averaged results from two technical replicates are shown for each condition. (D) Chitin-independent transformation efficiencies in DS40M4 and NBRC 15631 reactions with varied outgrowth times following addition of tDNA. Strains were transformed with 50 ng linear *luxO::spec*$^R$ tDNA. Averaged results from two technical replicates are shown for each condition.

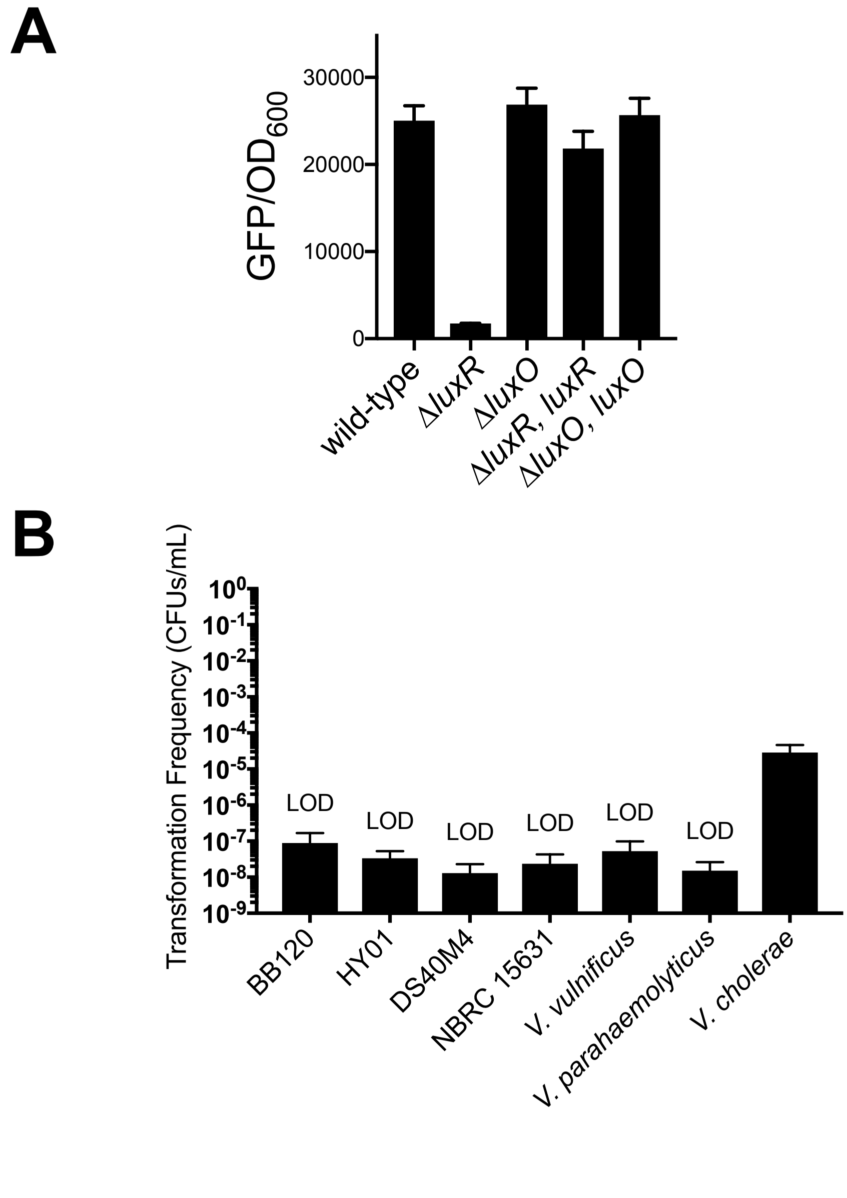


**Figure S2.** (A) Complementation of Δ*luxR* and Δ*luxO* in DS40M4. The $P_{luxCDABE}$-*gfp* reporter plasmid pCS019 was introduced into the strains of *V. campbellii,* and the GFP fluorescence divided by $OD_{600}$ was determined. (B) Chitin-dependent transformations in each of the listed *Vibrio* species using $Spec^R$ linear tDNAs (500 ng) targeting the *luxR* homolog in each strain. LOD, limit of detection.

**Table S1.** Strains used in this study.

| Strains | Genotype | Reference |
|---|---|---|
| ***V. campbellii* strains** | | |
| BB120 | wild-type (ATCC BAA-1116) | (8) |
| JAF78 | BB120 $\Delta luxO::CM^R$ | (9) |
| CAS156 | BB120 $\Delta luxO::CM^R$, pCS19 | This study |
| KM669 | $\Delta luxR$ | (10) |
| CAS094 | BB120 $\Delta luxR$, pCS19 | This study |
| CAS093 | BB120, pCS19 | This study |
| CAS190 | BB120, pCS32 | This study |
| CAS003 | BB120, pMMB67EH-tfox-kanR | This study |
| CAS165 | BB120, pMMB67EH-kanR | This study |
| DS4M04 | wild-type | (11) |
| CAS141 | DS40M4 $\Delta luxO::Spec^R$ | This study |
| CAS187 | DS40M4 $\Delta luxO::Spec^R$ $\Delta luxB::luxO::TM^R$ | This study |
| CAS189 | DS40M4 $\Delta luxO::Spec^R$ $\Delta luxB::luxO::TM^R$, pCS19 | This study |
| CAS179 | DS40M4 $\Delta luxO::Spec^R$ $\Delta luxB::luxO::TM^R$, pMMB67EH-tfoX-kanR | This study |
| CAS147 | DS40M4 $\Delta luxO::Spec^R$, pCS19 | This study |
| CAS043 | DS40M4 $\Delta luxO::Spec^R$, pMMB67EH-tfoX-kanR | This study |
| CAS211 | DS40M4 $\Delta luxR$ $\Delta luxB::TM^R$ | This study |
| CAS212 | DS40M4 $\Delta luxR$ $\Delta qstR::TM^R$ | This study |
| CAS096 | DS40M4 $\Delta luxR::Spec^R$ | This study |
| CAS105 | DS40M4 $\Delta luxR::Spec^R$, pCS19 | This study |
| CAS184 | DS40M4 $\Delta luxR::Spec^R$, pCS32 | This study |
| CAS095 | DS40M4 $\Delta luxR::Spec^R$, pMMB67EH-tfoX-kanR | This study |
| CAS186 | DS40M4 $\Delta luxR::Spec^R$, $\Delta luxB::luxR::TM^R$ | This study |
| CAS188 | DS40M4 $\Delta luxR::Spec^R$, $\Delta luxB::luxR::TM^R$, pCS19 | This study |
| CAS178 | DS40M4 $\Delta luxR::Spec^R$, $\Delta luxB::luxR::TM^R$, pMMB67EH-tfoX-kanR | This study |
| CAS181 | DS40M4 $\Delta qstR::TM^R$, pMMB67EH-tfoX-kanR | This study |
| CAS104 | DS40M4, pCS19 | This study |
| CAS213 | DS40M4, pCS32 | This study |
| CAS166 | DS40M4, pMMB67EH-kanR | This study |
| CAS034 | DS40M4, pMMB67EH-tfoX-kanR | This study |
| HY01 | wild-type | (12) |
| CAS106 | HY01, pCS19 | This study |
| CAS209 | HY01, pCS32 | This study |
| CAS168 | HY01, pMMB67EH-kanR | This study |
| CAS071 | HY01, pMMB67EH-tfoX-kanR | This study |
| NBRC 15631 | wild-type (CAIM 519, ATCC 25920) | (13) |
| CAS174 | NBRC 15631 $\Delta luxB::TM^R$ | This study |
| CAS142 | NBRC 15631 $\Delta luxO::Spec^R$ | This study |
| CAS148 | NBRC 15631 $\Delta luxO::Spec^R$, pCS19 | This study |

| | | |
|---|---|---|
| CAS045 | NBRC 15631 Δ*luxO::Spec*$^R$, pMMB67EH-tfoX-kanR | This study |
| CAS075 | NBRC 15631 Δ*luxR::Spec*$^R$ | This study |
| CAS103 | NBRC 15631 Δ*luxR::Spec*$^R$, pCS19 | This study |
| CAS217 | NBRC 15631 Δ*luxR::Spec*$^R$, pCS32 | This study |
| CAS044 | NBRC 15631 Δ*luxR::Spec*$^R$, pMMB67EH-tfoX-kanR | This study |
| CAS102 | NBRC 15631, pCS19 | This study |
| CAS210 | NBRC 15631, pCS32 | This study |
| CAS035 | NBRC 15631, pMMB67EH-tfoX-kanR | This study |
| CAS169 | NBRC15631, pMMB67EH-kanR | This study |
| ***V. cholerae* strains** | | |
| E7946 | wild-type 01 El Tor | (14) |
| SAD793 | E7946 Sm$^R$ Δ*hapR::Spec*$^R$, pMMB67EH-tfox-kanR | This study |
| CAS214 | E7946, pCS32 | This study |
| VC20 | E7946, pMMB67EH-tfoX-kanR | This study |
| CAS172 | E7946 Sm$^R$ Δ*hapR::Spec*$^R$, pMMB67EH-tfoX-kanR | This study |
| CAS177<br>VC21 | E7946, pMMB67EH-kanR<br>E7946 Sm$^R$ Δ*luxO::Spec*$^R$, pMMB67EH-tfox-kanR | This study<br>This study |
| ***V. natriegens strains*** | | |
| ATCC 14048 | wild-type (NBRC 15631) | (15) |
| CAS215 | wild-type (NBRC 15636), pCS32 | This study |
| CAS039 | Δ*dns::Spec*$^R$, pMMB67EH-tfoX-carbR | (15) |
| CAS001 | ATCC 14048, pMMB67EH-tfoX-kanR | This study |
| CAS036 | ATCC 14048, pMMB67EH-carbR | (15) |
| ***V. vulnificus strains*** | | |
| ATCC 27562 | wild-type | ATCC |
| CAS-vv013 | ATCC 27562, pMMB67EH-tfoX-kanR | This study |
| CAS-vv015 | ATCC 27562 Δ*smcR::Spec*$^R$, pMMB67EH-tfoX-kanR | This study |
| CAS-vv022 | ATCC 27562, pCS32 | This study |
| ***V. parahaemolyticus strains*** | | |
| RIMD2210633 | wild-type | ATCC |
| CAS-V01 | RIMD2210633, pMMB67EH-tfoX-kanR | This study |
| CAS-V03 | RIMD2210633 Δ*opaR::Spec*$^R$, pMMB67EH-tfoX-kanR | This study |
| CAS-V16 | RIMD2210633, pCS32 | This study |
| ***E. coli* strains** | | |
| S17-1λpir | Wild-type, mating strain | (16) |

**Table S2.** Plasmids used in this study.

| Name | Description | Reference |
|---|---|---|
| pMMB67EH-tfoX | $P_{tac}$::*tfoX*; *amp*$^R$ | (15) |
| pMMB67EH | Empty vector control; *amp*$^R$ | (17) |
| pMMB67EH-tfoX-kanR k | Derivative of pMMB67EH-tfoX; *kan*$^R$ | This study |
| pMMB67EH-kanR | Empty vector control; *kan*$^R$ | This study |
| pCS19 | $P_{luxCDABE}$::*gfp*; *kan*$^R$; derivative of pMMB67EH-tfoX-kanR | This study |
| pJV298 | $P_{tac}$-*gfp*; *lacIq*; colE1 origin, *CM*$^R$ | This study |
| pCS32 | $P_{tac}$::*tfoX-qstR*; *kan*$^R$; derivative of pMMB67EH-tfoX-kanR | This study |

**Table S3.** Oligos used in this study.

| Primer Name | primer sequence | Description |
|---|---|---|
| **Primers for mutant constructs** | | |
| lab179 | attccggggatccgtcgac | Ampflify Middle AbR (SpecR, TmR) F |
| lab180 | tgtaggctggagctgcttc | Ampflify Middle AbR (SpecR, TmR) R |
| CAS0001 | gctccgtcagaccactggcaacca | ΔLuxR BB120 F1 |
| CAS0002 | gtcgacggatccccggaatgtccatatttcttttt ccttgccatttgagttg | ΔLuxR BB120 R1 |
| CAS0003 | gaagcagctccagcctacactatgcatctac aaccgtgaacatcactaaaaaataattag | ΔLuxR BB120 F1 |
| CAS0004 | cgcatctccagttgttcgtcgtct | ΔLuxR BB120 R2 |
| CAS0069 | gtaccaccaatgccagaatcagtgc | NBRC 15361 ΔLuxR F1 |
| CAS0084 | ccacgcagctccttgtccgaatcg | DS40M4 ΔLuxR F1 |
| CAS0070 | gtcgacggatccccggaataggtctctttgca attgagtccat | NBRC 15361, DS40M4 ΔLuxR R1 |
| CAS0071 | gaagcagctccagcctacaatctacaaccgt gaacatcactaa | NBRC 15361, DS40M4 ΔLuxR F2 |
| CAS0072 | cgactacgcacacgcatctccag | NBRC 15361,DS40M4 ΔLuxR R2 |
| CAS0074 | gtccgtcgctctaccaactgagc | NBRC 15361, DS40M4 ΔLuxO F1 |
| CAS0075 | gtcgacggatccccggaattaccattagtag ataacgagac | NBRC 15361, DS40M4 ΔLuxO R1 |
| CAS0076 | gaagcagctccagcctacagtatgaatacg gacgtattaaatcagc | NBRC 15361, DS40M4 ΔLuxO F2 |
| CAS0077 | gcttgttcaactaggtagccaccaga | NBRC 15631 ΔLuxO R2 |
| CAS0079 | gtgcttctggcgtgctgtcacg | DS40M4 ΔLuxO R2 |
| CAS0101 | cgtgcttgttcacggctcaagc | HY01 ΔLuxR F1 |
| CAS0102 | gtcgacggatccccggaatctttgcaattgag tccataatcc | HY01 ΔLuxR R1 |
| CAS0103 | gaagcagctccagcctacagatatgctatgc atctacaaccgt | HY01 ΔLuxR F2 |
| CAS0104 | gctcctgcagcagaagcggctc | HY01 ΔLuxR R2 |
| CAS0107 | cgtctcaggcaacgcagacagtg | ATCC 27562 ΔSmcR F1 |

| | | |
|---|---|---|
| CAS0108 | gtcgacggatccccggaattgagtccatagg ttgtttccttacc | ATCC 27562 ΔSmcR R1 |
| CAS0109 | gaagcagctccagcctacagaacacgaata gcaccagtaacctc | ATCC 27562 ΔSmcR F2 |
| CAS0110 | ggaagctcaagcaacgactagtg | ATCC 27562 ΔSmcR R2 |
| CAS0113 | cgatacctgctagcactgccattg | RIMD2210633 ΔOpaR F1 |
| CAS0114 | gtcgacggatccccggaaaggtctctttgcaa ttgagtccatatcc | RIMD2210633 ΔOpaR R1 |
| CAS0115 | gaagcagctccagcctacacgcgaacacta aagctcagatttg | RIMD2210633 ΔOpaR F2 |
| CAS0116 | gacactggcatgaagatcactccac | RIMD2210633 ΔOpaR R2 |
| CAS0134 | gcagctcgcgatccgaatcatgct | N16961 ΔHapR F1 |
| CAS0135 | ggcttcaaccacacgttcaccat | N16961 ΔHapR R2 |
| CAS0148 | cgtgctcaagtcttcactgatgatg | DS40M4 and NBRC ΔluxB F1 |
| CAS0149 | gtcgacggatccccggaatgatgacttgatc agaagaacgctttga | DS40M4 ΔluxB R1 |
| CAS0150 | gaagcagctccagcctacacactcgtaacgt ttaaacgatgctgag | DS40M4 ΔluxB F2 |
| CAS0151 | ggtgaatggccacaaggtacct | DS40M4 and NBRC ΔluxB R2 |
| CAS0206 | gtcgacggatccccggaatgaagaataatc caaatttcatgtctc | R1 delta LuxB NBRC |
| CAS0207 | gaagcagctccagcctacagtcaaatacca ctcgtaacgtttaaac | F2 delta LuxB NBRC |
| BBC1264 | in ad list | F1 Δdns V. nat |
| BBC1267 | in ad list | R2 Δdns V. nat |
| CAS0247 | ctaggtagatactgctcttctggagag | F1 to delete QstR in DS40M4 |
| CAS0248 | gtcgacggatccccggaatagcatcctcttcc atgctgattag | R1 to delete QstR in DS40M4 |
| CAS0249 | gaagcagctccagcctacactgatgtcataa aacaatgatgagcaac | F2 to delete QstR in DS40M4 |
| CAS0250 | caactgaacaagccaacaggaacg | R2 to delete QstR in DS40M4 |
| CAS0252 | gaagcagctccagcctacaataaagtcgact tggtgagtcagtc | F to amplify LuxR to complement into DS4 with homology to Tm cassette |
| CAS0253 | tcgtttaaacgttacgagtgttagtgatgttcac ggttgtagatgc | R to amplify LuxR to complement into DS4 with homology to down stream of LuxB |
| CAS0254 | cactcgtaacgtttaaacgatgctg | F2 to amplify down arm of 4 piece SOE product for complementing LuxR or LuxO, by deleting LuxB and replacing with TmR and LuxR or LuxO gene |
| CAS0255 | gaagcagctccagcctacacaacagttgga gaaggagatcagtc | F to amplify LuxO to complement into DS4 with homology to Tm cassette |
| CAS0256 | tcgtttaaacgttacgagtgcatacgttttgtttt cgtccttgc | R to amplify LuxO to complement into DS4 with homology to down stream of LuxB |
| CAS0295 | gctaattcagtttaagcggccataggtctctttg caattgagtccat | R1 to make delta LuxR unmarked in DS40M4 |
| CAS0296 | atggccgcttaaactgaattagcatctacaac cgtgaacatcactaa | F2 to make delta LuxR unmarked in DS40M4 |
| CAS0334 | cacgagcaagatggttgttaagc | F1 V. vulnificus delta pomB(motB) Tm SOE |
| CAS0335 | gtcgacggatccccggaatcatcacatactc ccgtgattaatcattg | R1 V. vulnificus delta pomB(motB) Tm SOE |

| | | |
|---|---|---|
| CAS0336 | gaagcagctccagcctacagagcagtaatt gggtacgtgagttg | F2 V. vulnificus delta pomB(motB) Tm SOE |
| CAS0337 | cacgtcaatgtctggctctttagc | R2 V. vulnificus delta pomB(motB) Tm SOE |
| CAS0339 | gatgttcttcagtgcgacgaaccag | F1 V.para delta pomB(motB) Tm SOE |
| CAS0340 | gtcgacggatccccggaatcatcacaaatct ccgcgattactc | R1 V.para delta pomB(motB) Tm SOE |
| CAS0341 | gaagcagctccagcctacagttattcaataac aaagcgcgtc | F2 V.para delta pomB(motB) Tm SOE |
| CAS0342 | gatcgctacatctacatcatcagtca | R2 V.para delta pomB(motB) Tm SOE |
| **Detection primers for mutant constructs** | | |
| CAS0083 | gaagcagctccagcctaca | F detect for all AbR Cassettes |
| CAS0073 | gtgatgcagaagatatcgcac | R detect for NBRC ΔLuxR |
| CAS0078 | tgtcgatggcaatcgctagttcac | R detect for NBRC ΔLuxO |
| CAS0080 | cgtagtcacggagtcattggcttc | R detect for DS40M4 ΔLuxO |
| CAS0085 | gaaggctcaatcactgaccttcc | R detect for DS40M4 ΔLuxR |
| CAS0094 | gttcttgttgctatcggtcgtgttcc | R detect for BB120 ΔLuxR |
| CAS0112 | gcagctccagtagctgcacctg | R detect for ATCC 27562 ΔSmcR |
| CAS0118 | cgtcgtcacttcgaagtgaacg | R detect for RIMD2210633 ΔOpaR |
| CAS0220 | gctgagcatcaatcgctcttgac | DS40M4 ΔluxB detect |
| CAS0251 | ggttcatcacctacaagctcacgac | detect primer delta QstR in DS4 |
| CAS0338 | gagcacttgttggtaatattgatggc | R detect vulnificus delta pomB(motB) Tm SOE |
| CAS0343 | gttgctctcagaggtcacttcaatg | R detect V.para delta pomB(motB) Tm SOE |
| CAS0152 | atggccgcttaaactgaattagc | MASC-PRC Forward detect |
| **Primers for plasmids** | | |
| CAS0125 | agcttggctgttttggcgga | pMMB tfoX F |
| CAS0126 | ggcctatggagctgtgcggc | pMMB tfoX R |
| CAS0127 | actgagcgctgccgcacagctccataggcc cttatgaagtccatactttttcactga | PluxC-GFP + homology to pMMB, F |
| CAS0129 | tcttctctcatccgccaaaacagccaagcttc agttgtacagttcatccatgc | PluxC-GFP + homology to pMMB, R |
| CAS0289 | caggatcccgggaggaggtaggcgcgtga aaaaatcggcttatgcgag | F to amplify QstR DS40M4 insert with homology to pMMB-tfox, includes RBS |
| CAS0290 | tccgccaaaacagccaagctttatgacatca ggttttggtcag | R to amplify QstR DS40M4 insert with homology to pMMB-tfox, |
| CAS0293 | cctcctcccgggatcctgtgtgaaattgaggtc gactctagaggatccttaac | R to amplify the pmmb-tfox backbone to insert QstR |
| CAS0294 | agcttggctgttttggcggatg | F to amplify the pmmb-tfox backbone to insert QstR |

**Table S4.** BB120 and DS40M4 homologs of *V. cholerae* competence genes.

| *V. cholerae* locus tag | DS40M4 locus tag | BB120 locus tag | Gene name | Competence category |
|---|---|---|---|---|
| VC0611 | DSB67_12495 | VIBHAR_03432 | | chitin |
| VC0612 | DSB67_12490 | VIBHAR_03431 | | chitin |
| VC0613 | DSB67_12485 | VIBHAR_03430 | | chitin |
| VC0614 | DSB67_12480 | VIBHAR_03429 | | chitin |
| VC0615 | DSB67_12475 | VIBHAR_03428 | | chitin |
| VC0616 | DSB67_12470 | VIBHAR_03427 | | chitin |
| VC0617 | DSB67_12465 | VIBHAR_03426 | | chitin |
| VC0618 | DSB67_12460 | VIBHAR_03425 | | chitin |
| VC0619 | DSB67_12455 | VIBHAR_03424 | | chitin |
| VC0620 | DSB67_12450 | VIBHAR_03423 | *cbp* | chitin |
| VC0622 | DSB67_12445 | VIBHAR_03422 | *chiS* | chitin |
| VC0972 | DSB67_03685 | VIBHAR_01269 | chitoporin | chitin |
| VC0995 | DSB67_03995 | VIBHAR_01336 | glcNAcPTS | chitin |
| VC1952 | DSB67_11840 | VIBHAR_03258 | *chiA1* | chitin |
| VC2080 | DSB67_04120 | VIBHAR_01362 | *tfoS* | chitin |
| VCA0027 | DSB67_19440 | VIBHAR_06955 | *chiA2* | chitin |
| VCA0700 | DSB67_23890 | VIBHAR_05945 | chitodextrinase | chitin |
| VC0470 | DSB67_13165 | VIBHAR_03571 | *dns* | inhibition |
| VC0766 | DSB67_02900 | VIBHAR_01075 | *exoVII* | inhibition |
| VC2417 | DSB67_02380 | VIBHAR_00958 | *recJ* | inhibition |
| VC0032 | DSB67_15695 | VIBHAR_00409 | *comM* | integration |
| VC0048 | DSB67_15605 | VIBHAR_00389 | *dprA* | integration |
| VC0543 | DSB67_12880 | VIBHAR_03513 | *recA* | integration |
| VC1879 | DSB67_04755 | VIBHAR_01534 | *comEC* | integration |
| VC1917 | DSB67_04445 | VIBHAR_01422 | *comEA* | integration |
| VC2719 | DSB67_00645 | VIBHAR_00615 | *comF* | integration |
| VC0462 | DSB67_13195 | VIBHAR_03577 | *pilT* | pilus |
| VC0857 | DSB67_03055 | VIBHAR_01136 | | pilus |
| VC0858 | DSB67_03060 | VIBHAR_01137 | | pilus |
| VC0859 | DSB67_03065 | VIBHAR_01138 | | pilus |
| VC0860 | DSB67_03070 | VIBHAR_01139 | | pilus |
| VC0861 | DSB67_03075 | VIBHAR_01140 | | pilus |
| VC1612 | DSB67_08830 | VIBHAR_02493 | | pilus |
| VC2423 | DSB67_12665 | VIBHAR_03468 | *pilA* | pilus |
| VC2424 | DSB67_12670 | VIBHAR_03469 | *pilB* | pilus |
| VC2425 | DSB67_12675 | VIBHAR_03470 | *pilC* | pilus |
| VC2426 | DSB67_12680 | VIBHAR_03471 | *pilD* | pilus |

| | | | | |
|---|---|---|---|---|
| VC2630 | DSB67_13985 | VIBHAR_00029 | *pilQ* | pilus |
| VC2631 | DSB67_13990 | VIBHAR_00030 | *pilP* | pilus |
| VC2632 | DSB67_13995 | VIBHAR_00031 | *pilO* | pilus |
| VC2633 | DSB67_14000 | VIBHAR_00032 | *pilN* | pilus |
| VC2634 | DSB67_14005 | VIBHAR_00033 | *pilM* | pilus |
| VC0396 | DSB67_13770 | VIBHAR_03706 | *qstR* | regulation |
| VC0534 | DSB67_12895 | VIBHAR_03517 | *rpoS* | regulation |
| VC0583 | DSB67_12620 | VIBHAR_03459 | *hapR* | regulation |
| VC1153 | DSB67_06270 | VIBHAR_02628 | *tfoX* | regulation |
| VC2677 | DSB67_01305 | VIBHAR_00726 | *cytR* | regulation |
